# iVirP: An integrative, efficient, and user-friendly pipeline to annotate viral contigs from raw reads of metagenome or VLP sequencing

**DOI:** 10.1101/2024.01.21.576577

**Authors:** Bowen Li, Xianyue Jiao, Guanxiang Liang

## Abstract

Metagenome sequencing and virus-like particles sequencing make it possible to explore the virome in the humans and other organisms. One way to analyze the sequencing data is to assemble reads into contigs according to the overlapping regions, and then the predicted viral contigs are screened out to conduct deeper dives. iVirP (integrative virome pipeline) is a user-friendly pipeline that includes the whole process of viral contigs discovery from the quality control of raw data to the filter of high quality viral contigs. This pipeline also features a branching function that can estimate the abundance of known eukaryotic viruses in a short period, while reducing host contamination. It is suitable for the rapid diagnosis of pathogens. Throughout iVirP, many details that might affect the experience of users are optimized carefully to reduce the time spent on dealing with usage and errors. iVirP was tested on a published, high-quality VLP sequencing dataset and was able to well reproduce the conclusions of the corresponding research. The benchmark indicates that iVirP could accurately assemble viral contigs from real sequencing data. iVirP is easy to install and currently available at https://github.com/li-bw18/iVirP.

## 1. Introduction

The metagenomic sequencing and virus-like particles (VLPs) sequencing are always performed to explore the landscape of the virome in the humans and other animals (Harvey and Holmes 2022; Liang and Bushman 2021). To process and analyze the sequencing data, one way is to align reads to the reference viral genomes (Czeczko et al. 2017). However, there are still many unknown or novel viruses that have not been included in the reference. Therefore, assembly-based methods, which assemble the putative genomes of viruses according to the overlapping sequences between different reads in single or multiple libraries, are widely utilized these years with the rapid progression of high-performance computers. The genomes assembled in this process are referred to as viral contigs. After viral contigs are identified using assembly-based methods, various downstream analyses can be conducted to investigate both known and unknown viruses (Liang et al. 2020; Norman et al. 2015).

Obtaining high-quality viral contigs from raw data requires several sequential steps. To enhance the efficiency and convenience of these processes, the imperative development of an integrative pipeline for comprehensive analysis is warranted. There exist some related pipelines, for example, VIRify (Rangel-Pineros et al. 2023) and ViWrap (Zhou et al. 2023) could identify viral contigs from the assembled contigs and make some annotations on these viral contigs, while ViroProfiler (Ru J et al. 2023) could both identify viral contigs from raw reads and accomplish some downstream analysis. However, focusing too much on the comprehensiveness of functions will make a pipeline difficult to install and run, so for the user-friendliness, we developed a novel pipeline iVirP, which only includes the routine upstream virome analysis. iVirP could complete the whole process of viral contigs identification with a single command and limited essential parameters. The default values for the remaining parameters were carefully selected by the research team, and a benchmark assessment substantiated their appropriateness and acceptability. iVirP accepts the raw sequencing files as the input and outputs a sequence file of the discovered viral contigs and some other information files. In addition, as an attempt to rapidly diagnose clinical pathogens, we developed a branching function to estimate the abundance of some eukaryotic viruses in a sample with alignment.

## 2. Materials and Methods

### 2.1 The overview of iVirP

The architecture of iVirP is shown in **Fig. 1**. The main line of iVirP could generally be divided into three parts according to the color of the boxes. The very first part colored yellow is the quality control steps, in which the reads with sequences of adapters and animal hosts are trimmed with Trimmomatic (Bolger et al. 2014) and Bowtie2 (Langmead and Salzberg 2012), and meanwhile, two quality reports are given by FastQC (Andrews 2010) and ViromeQC (Zolfo et al. 2019), respectively. In the second part colored blue, the cleaned reads are assembled into contigs with SPAdes (Prjibelski et al. 2020). Finally, in the last part colored red, contigs shorter than 1000bp are discarded with VSEARCH (Rognes et al. 2016), and among the remaining contigs, those who meet at least one of the following two requirements are defined as potential viral contigs. The first is that a contig could be significantly aligned to any of the viral reference databases selected by us with BLASTN (Camacho et al. 2009), while the second is that a contig is predicted as a viral sequence by VirSorter2 (Guo et al. 2021), which is based on the machine learning. Then the potential viral contigs are further filtered with CheckV (Nayfach et al. 2021a), retaining those with high quality and completeness as the final viral contigs. Before the final output, these viral contigs are clustered with VSEARCH to obtain a non-redundant set. The different kinds of information about these non-redundant viral contigs during filtering, including whether the contigs are identified by BLASTN or VirSorter2, which genome of which reference database is aligned by each viral contig, and the detailed quality reports from CheckV, are all output in the form of tables for the downstream analysis.

**Figure 1.**
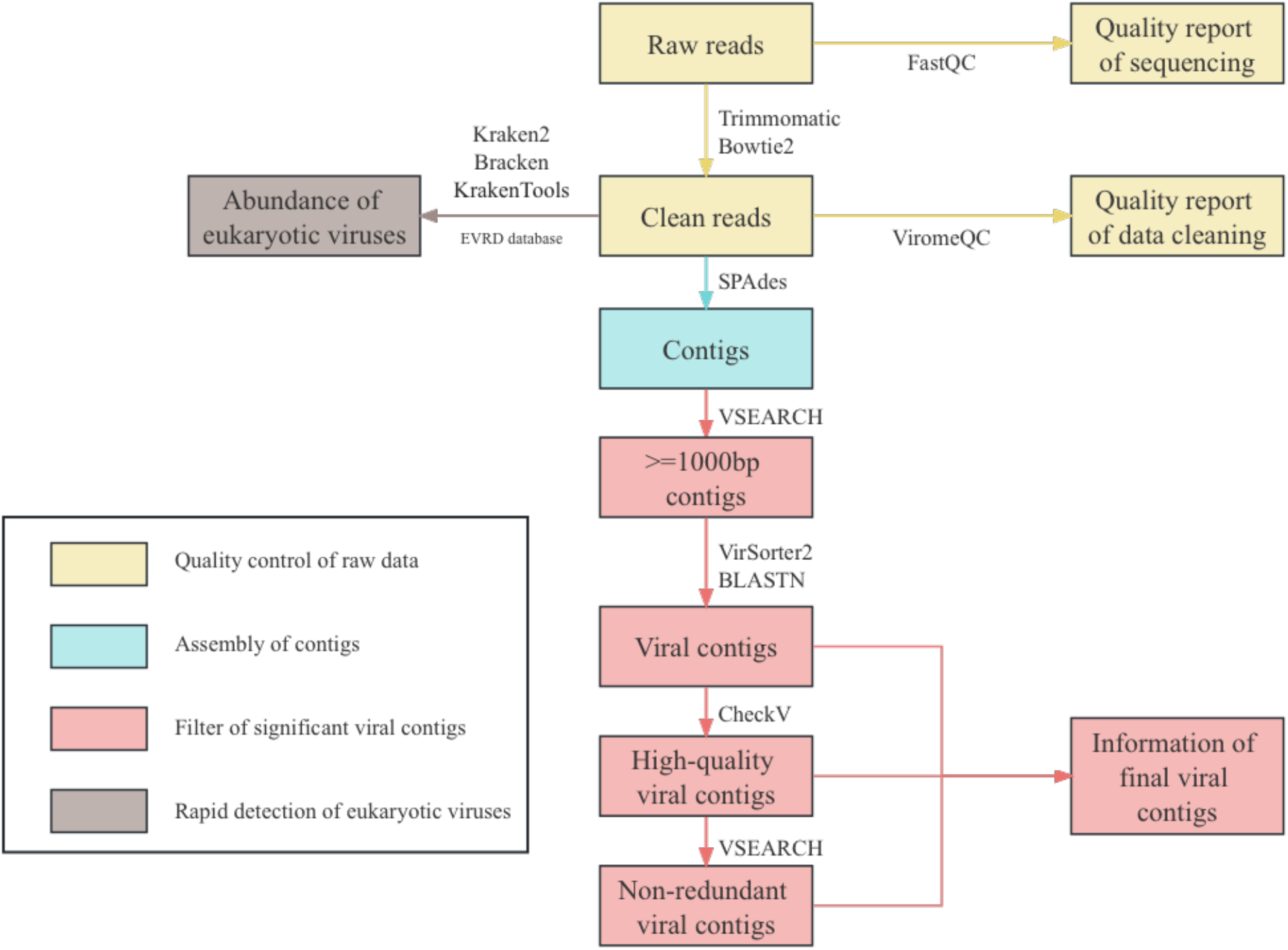
The architecture of iVirP. The main line of iVirP consists of three modules, as well as a branching function module, each of which is mentioned in the legend. The software used, the inputs, and the outputs of each step are included in the texts of the flowchart.

As for the branching line, the reads after the quality control steps are mapped to the EVRD database (Chen et al. 2022), a nonredundant and well-refined eukaryotic viral reference database, with Kraken2 (Wood et al. 2019) to obtain the estimated abundance of common eukaryotic viruses. Then Bracken2 (Lu et al. 2017) and KrakenTools (Lu et al. 2022) are utilized to get more accurate estimates and more visible outputs, respectively.

### 2.2 Detailed designs and parameters in iVirP

The parameters that are necessary for running the main line of iVirP are only the paths of the raw sequencing files. The adapters, the host reference sequences to be removed, the thread, and the output path are adjustable parameters with default values, and the rest of the parameters are not adjustable. These non-adjustable parameters are set as follows. The parameters of FastQC, Trimmomatic, Bowtie2, ViromeQC, SPAdes, and VirSorter2 are set to the default values of the corresponding software or to the consensus values for the corresponding analysis. When VSEARCH filters the contigs according to the length, the minimum value is set to 1000bp, and the maximum value is not set, while when VSEARCH performs the clustering of the final viral contigs, the similarity parameter is set to 0.995, and the fast cluster mode is chosen. The criteria for a significant alignment in BLASTN are “evalue” not higher than 10^−10^, “pident” not lower than 50, and “qcovs” not lower than 80. In CheckV, only viral contigs that are identified as “Complete”, “High-quality,” and “Medium-quality” will be retained. The parameters in the branching function are all default values of the corresponding software. The default settings of these non-adjustable parameters are based on some well-known literatures related to the analysis of virome data and our experiences (Kaelin et al. 2022; Shkoporov et al. 2019; Shkoporov et al. 2022).

The reference virus databases used for BLASTN alignment include NCBI Refseq, GPD (Camarillo-Guerrero et al. 2021), GVD (Gregory et al. 2020), MGV (Nayfach et al. 2021b), and crAss (Guerin et al. 2018), all of which are carefully selected. According to our experiences, these five databases as a whole have strong representation and coverage, enough for most situations.

### 2.3 User-friendly optimizations

iVirP has several optimizations in terms of user-friendliness. First of all, as mentioned earlier, most of the parameters in iVirP do not need to be adjusted. Although this makes some unusual sequencing raw files unfit for iVirP, it is acceptable compared to the benefits of concise parameters. The second is that iVirP provides two different installation methods, both of which have been debugged for many times on different computers, allowing users to install iVirP with the simplest commands and the highest probability of success. The third is that iVirP supports the breakpoint reconnection during both installation and running. In other words, if the progress is interrupted for some reasons when installing or running iVirP, users could continue the progress from the interrupted step after the problems are eliminated, reducing the time spent on exploring the software. Lastly, iVirP organizes intermediate and result files as neatly as possible in the output path and provides a lot of information that might be needed in the downstream analysis for users, reducing the possibility of re-running some steps for certain information.

## 3. Results

To demonstrate that the main line of iVirP can accurately and efficiently analyze the virome data, we used iVirP to perform a reproduction analysis of 60 VLP sequencing samples from 20 infants in a classical virome analysis work Liang et al. (2020), and the results show that the number of viral contigs obtained by iVirP is consistent with the original analysis (**Fig. 2**).

**Figure 2.**
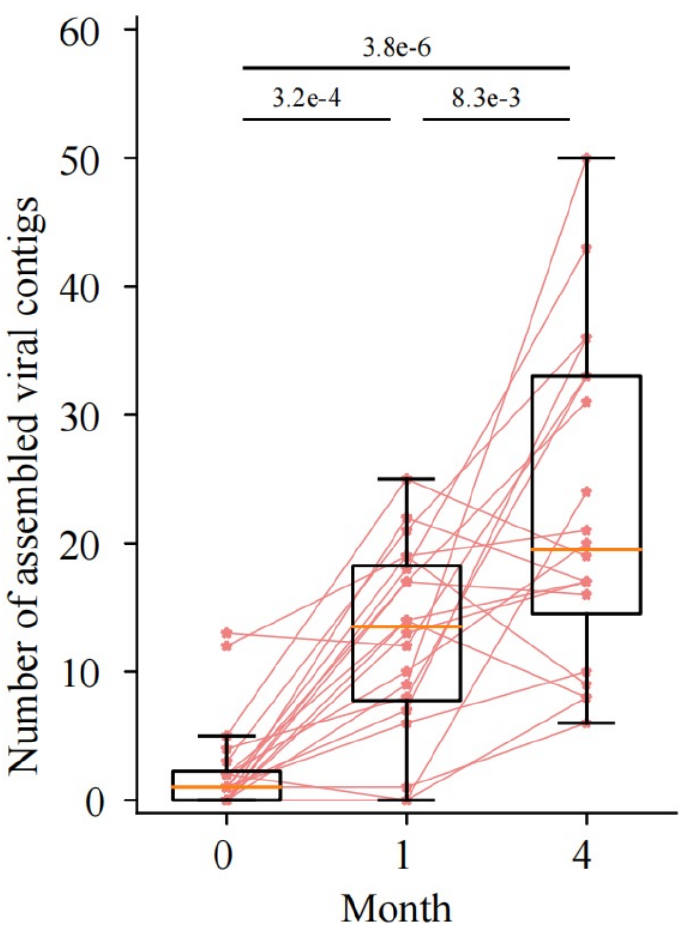
The number of assembled viral contigs in each sample of the reproduction analysis. Similar to the results of the original analysis, the number of putative viral genomes in the stool samples of the newborn, 1 month, and 4 months of age gradually increases.

The taxonomy of viral contigs identified by iVirP could be annotated utilizing the outcomes of BLASTN alignment, and the cleaned reads were aligned to these contigs using Bowtie2 to obtain the abundance of different virus families. Notably, the results are consistent with the initial analysis (**Fig. 3**). This benchmark proves that the main line of iVirP is capable of the assembly-based virome analysis.

**Figure 3.**
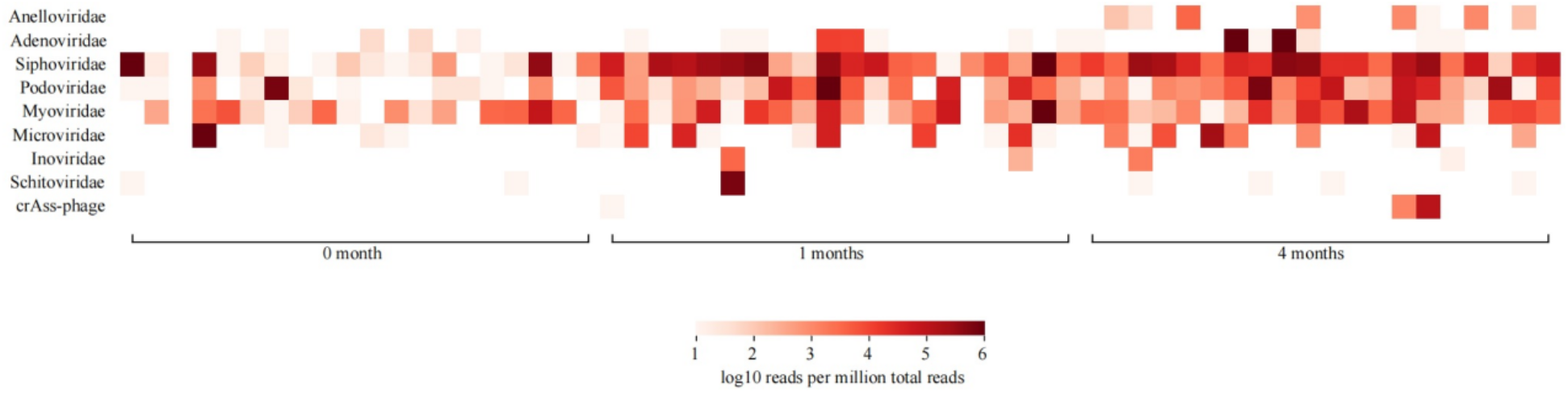
The abundance of different virus families in each sample of the reproduction analysis. Although the included family categories are not exactly the same as those in the original analysis (possibly due to the differences in taxonomy annotation methods and databases), the overall trend of the heatmap is largely consistent.

## 4. Discussions

The main line of iVirP is now relatively complete, but sacrificing the freedom to adjust parameters for user-friendliness might be an important limitation, so in the future, we might try to add some adjustable parameters without significantly increasing the burden on users, allowing iVirP to be fit for more situations. At the same time, it is also possible to add some commonly used downstream analysis to iVirP, such as taxonomy annotation, statistical figures plotting, contigs abundance calculation, host prediction, and even gene-level analysis, although this might make it more difficult to configure the virtual Python environment for iVirP. In the long term, we will also consider developing algorithms to improve the performance of some steps in the pipeline for discovering virus contigs with better quality.

We are also considering the improvement of the branching function, which might mainly rely on the algorithm design to speed up the estimation with as few false positives as possible, to better apply it to the clinical scenarios.

